# Acoustic Ejection Mass Spectrometry for High-Throughput Analysis

**DOI:** 10.1101/2020.01.28.923938

**Authors:** Hui Zhang, Chang Liu, Wenyi Hua, Lucien P. Ghislain, Jianhua Liu, Lisa Aschenbrenner, Stephen Noell, Kenneth Dirico, Lorraine F. Lanyon, Claire M. Steppan, Don W. Arnold, Thomas R. Covey, Sammy S. Datwani, Matthew D. Troutman

## Abstract

We describe a mass spectrometry (MS) analytical platform resulting from the novel integration of acoustic droplet ejection (ADE) technology, an open-port interface (OPI), and electrospray ionization (ESI) MS that creates a transformative system enabling high-speed sampling and label-free analysis. The ADE technology delivers nanoliter droplets in a touchless manner with high speed, precision and accuracy; subsequent sample dilution within the OPI, in concert with the capabilities of modern ESI-MS, eliminates the laborious sample preparation and method development required in current approaches. This platform is applied to a variety of experiments, including high-throughput (HT) pharmacology screening, label-free in situ enzyme kinetics, in vitro and in vivo adsorption, distribution, metabolism, elimination, pharmacokinetic (PK) and biomarker analysis, and HT parallel medicinal chemistry.

**One Sentence Summary:** ADE-OPI-MS is a transformational analytical platform that increases mass spectrometry utility via sub-second speed and non-contact sampling.

---

Mass is a fundamental molecular characteristic, and the advent of mass spectrometry (MS) to measure it has had a profound impact on science. In recent times, mass spectrometers have become commonplace due to their commercially availability, ruggedness and user-friendly interfaces, while offering exquisite capabilities. Modern instruments can routinely measure mass with high accuracy (>PPM) and high speed (<0.1 s) despite requiring miniscule sample volumes (<nL) (*1, 2*). A key advantage of MS is its ability to reliably measure a wide variety of analytes, from small molecules to proteins. Furthermore, many mass spectrometers allow gas-phase experiments within the instrument, offering near-certain analytical fidelity. Taken together, it is easy to see how these instruments have become core technologies when high-fidelity, high-sensitivity analysis is critical to enabling scientific progression, decision-making and supporting regulatory documentation.

While MS has made tremendous strides, there remain limitations. Effective and reproducible sample delivery to the instrument is a key element in successful experimental MS analysis. For aqueous samples, liquid chromatography (LC), following some level of sample workup, is typically employed for sample introduction. While this approach has proven effective, it is not entirely complementary to MS, as LC and MS are mis-matched with regard to throughput and sample requirements. This is not a limitation in cases such as untargeted analytical experiments, where a broad range of analytes need to be profiled (e.g., proteomics). However, for targeted analysis, use of LC greatly limits MS throughput and performance. This is a function of both the time required (for sample preparation and analysis) and, perhaps more importantly, complexity due to the expert knowledge required to effectively maintain and optimize the LC aspect of the experiment. Providing a faster, simpler alternative to LC would significantly expand the impact and application of MS to additional measurement-based sciences.

Bioanalytical technologies typically utilized for analysis of liquid samples include radiometric approaches (*3*), optical approaches such as fluorescence spectroscopy (*4, 5*), and mass-based approaches including LC-MS (*6–8*), matrix-assisted laser desorption/ionization MS (MALDI-MS) (*9–11*), and solid-phase extraction MS (SPE-MS) (*12–14*). With regard to acoustic-based sample delivery, early attempts have focused on acoustic mist ionization (AMI-MS) approaches (*15, 16*). AMI-MS utilizes an acoustic transducer to create a cloud of charged femtoliter-volume droplets via pulsed ejection with ambient ionization and direct introduction to the MS via a heated transfer tube. This approach could be tractable, but routine application will require that key limitations and hurdles be surmounted.

In this work, we describe a novel sample delivery approach that will revolutionize how MS is accessed and utilized. The approach combines technologies that are synergistic for MS performance: acoustic droplet ejection (ADE) and an open-port interface (OPI) (*17*). The ADE technology can sample nL volumes accurately and precisely on the hertz timescale and, notably, the ADE transducer renders samples directly, such that no workup is required. The OPI provides an elegant, minimalist approach for sample introduction into the MS. With OPI, solvent flows continuously through a co-axial tube at a constant rate, allowing samples to be diluted online. In the case of a nL sample from ADE, it is trivial to achieve >1000-fold dilutions; this feature nearly eliminates the ion suppression that must be controlled for successful ESI-MS analyses (a key driver for the use of LC in conventional approaches). The ADE-OPI touchless transfer approach minimizes carry-over artifacts that can confound LC methods, as samples are introduced acoustically, followed by high-fold dilution with OPI solvent flow. Furthermore, optimizing the ADE-OPI element of the analysis is facile; hence, sample analysis using ESI-MS is greatly simplified, as well as more robust. In this report, we describe the instrument in some detail and characterize its fundamental analytical performance. Additionally, we highlight its performance in a variety of applications, from analyzing chemical reactions to in vivo and in vitro biological analytical quantification, by comparing conventional and ADE-OPI-MS approaches. The demonstrated gains in performance and utility should lead to a transformational expansion of MS as the readout of choice for quantitative measurement-based sciences.

The OPI has shown great potential as a robust sample delivery method for ESI-MS (*18*) analysis of multiple types of complex samples, including polymers, inks, oils, and tissues, (*17, 19, 20-22*). Further, OPI was utilized as the direct sampling interface to ESI-MS solid-phase micro-extraction (SPME) (*20, 21*). The OPI uses vertically aligned, co-axial tubes to deliver solvent through the tubing annulus to the capture port, where a vortex is formed. For the ADE-OPI-MS approach, we integrate ADE technology with an OPI, such that the droplets are delivered to a single position (an inverted OPI) and fluid transport introduces the sample to the detector (Fig. 1A and S1). More specifically, the inverted OPI is aligned above the source well of a microplate to capture nanoliter ADE samples. Here, the droplet volume is precisely controlled, independent of the sample matrix composition (Fig. S2). Peak width in the subsequent mass-response plot is a function of droplet dilution, dispersion and mixing, which spread the sample inside the transfer capillary as it is convectively transported to the ESI source (Fig. 1A). Dispersion in the transfer capillary typically limits the sampling rate, as the sample peaks diffuse into the carrier solvent and broaden over the transport time, eventually merging. Fig.1B demonstrates typical analytical traces obtained with a sampling rate of 2 Hz [baseline peak width 0.5 seconds, 2.5% CV. Higher sampling rates are possible with multiplexing (Fig. S3)]. At this speed, ADE-OPI-MS performance allows for a sample throughput of 50,000 to 100,000 samples per day, based on a standard 16-hour unattended run time and analysis speed (1-2 Hz). The use of ESI-MS allows for excellent quantification performance in terms of sensitivity, reproducibility, and linear dynamic range without carry-over, and is applicable to a wide range of analytes (Fig. S4-S6). Taken together, it is clear that the ADE-OPI-MS approach provides dramatic increases in sample analysis while maintaining, or perhaps improving, the analytical performance of conventional approaches.

**Fig. 1.**
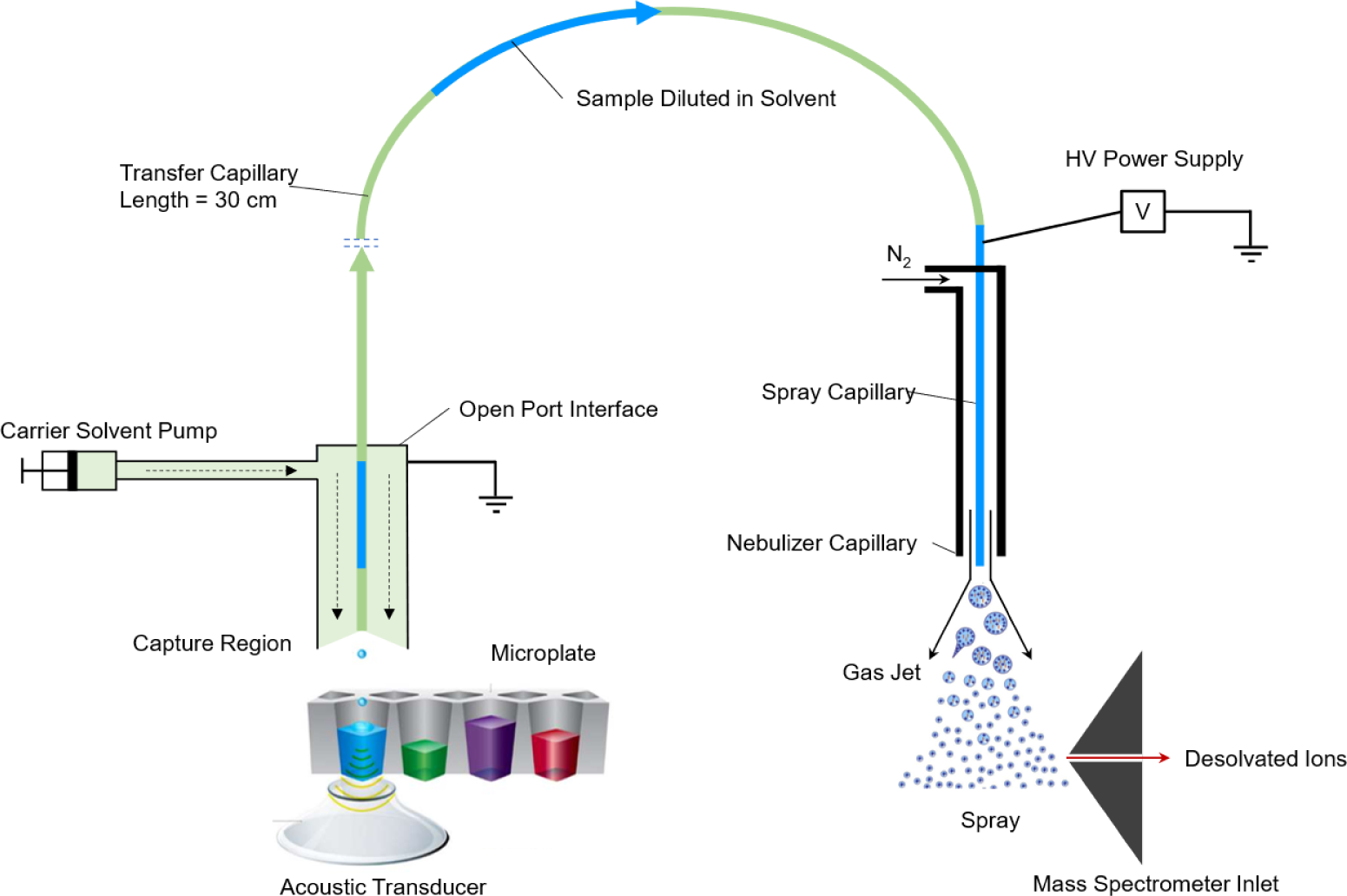
(A) Schematic of an ADE-OPI-MS system. A pulse of acoustic energy ejects sample droplets (2.5 nL) upwards at a velocity of 1 meter/second into the inverted OPI. A fluid pump delivers carrier solvent (300 μL/min) to a sample capture region that has a flow-stabilized vortex interface; sample is captured and diluted into a vortex of flowing carrier solvent. A high voltage (HV) supply and nebulizing gas (nitrogen) at the spray capillary drive conventional ESI.

**Fig. 1.**
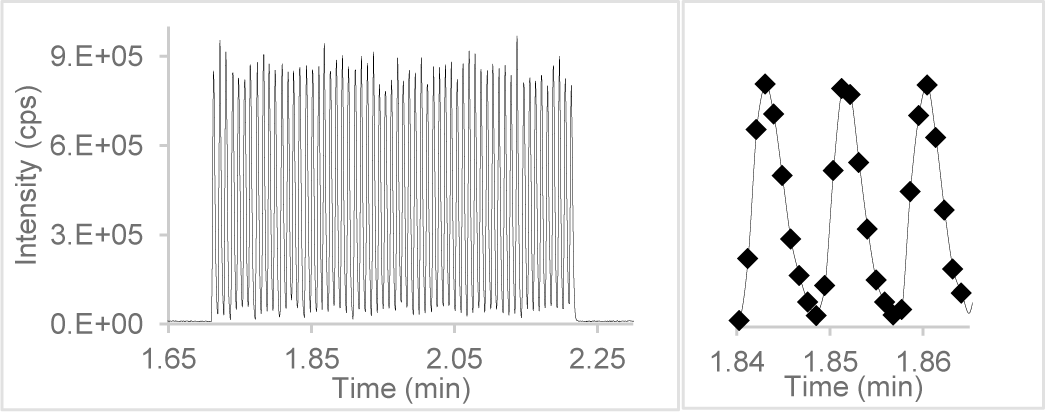
(B) Demonstration of 2 Hz sampling rate with neat acetonitrile as the carrier solvent (600 μL/min). Each injection consists of 5 nL of 100 nM dextromethorphan in water. Average full-width half-maximum (FWHM) is 0.21 sec, raw peak area CV is 2.5%.

To demonstrate the performance of ADE-OPI-MS as a quantitative reader for measurement-based approaches, we first assess the matrix tolerance with human plasma samples (Fig. 2). Plasma consists of a complex matrix that can dramatically impact analytical measurements; indeed, at present, direct analysis of plasma samples via ESI-MS is not possible. Typically, successful plasma sample analysis involves optimizing and performing time- and labor-intensive sample clean-up and LC separation steps. Using the ADE-OPI-MS approach, we tested sets of samples generated via a standard protein precipitation (Group 1) and 3 simple conditions (Groups 2, 3 and 4). Plasma samples prepared by simple 1:1 dilution with water and water containing 0.1% formic acid (v/v) (Groups 2 and 3, respectively) exhibited no significant difference when compared with those subjected to protein precipitation (Group 1) as regards symmetry of peak shape and sensitivity. Among these three groups, samples diluted with water containing 0.1% formic acid (v/v) (Group 2) provide optimal quantitation results (for accuracy and linearity of the standard curve). Neat plasma samples (e.g., untreated; Group 4) were also analyzed directly by ADE-OPI-MS; here, a moderate loss of signal (<2X) was observed, as compared to 1:1 diluted or protein-precipitated samples. For all treatment groups, these results were obtained without use of an internal standard and show that the ADE-OPI-MS platform is highly tolerant to complex matrices due to the nanoliter volume of sample injected, as well as the dilution effected within the OPI. [A demonstration of matrix tolerance with detergent is detailed in the supplementary information (Fig S5).] In this example, ADE-OPI-MS provides significant improvement versus conventional approaches in two dimensions: 1) simplicity and efficiency of operation via near-elimination of sample preparation and 2) speed of analysis. Another key advantage is the sample-sparing nature of the ADE-OPI-MS: the minimal sample volume required and avoidance of sample-destroying preparation steps allow for a reduction in plasma sampling volume with a corresponding preservation of samples such that original, unaltered samples may be re-analyzed repeatedly at later dates.

**Fig. 2.**
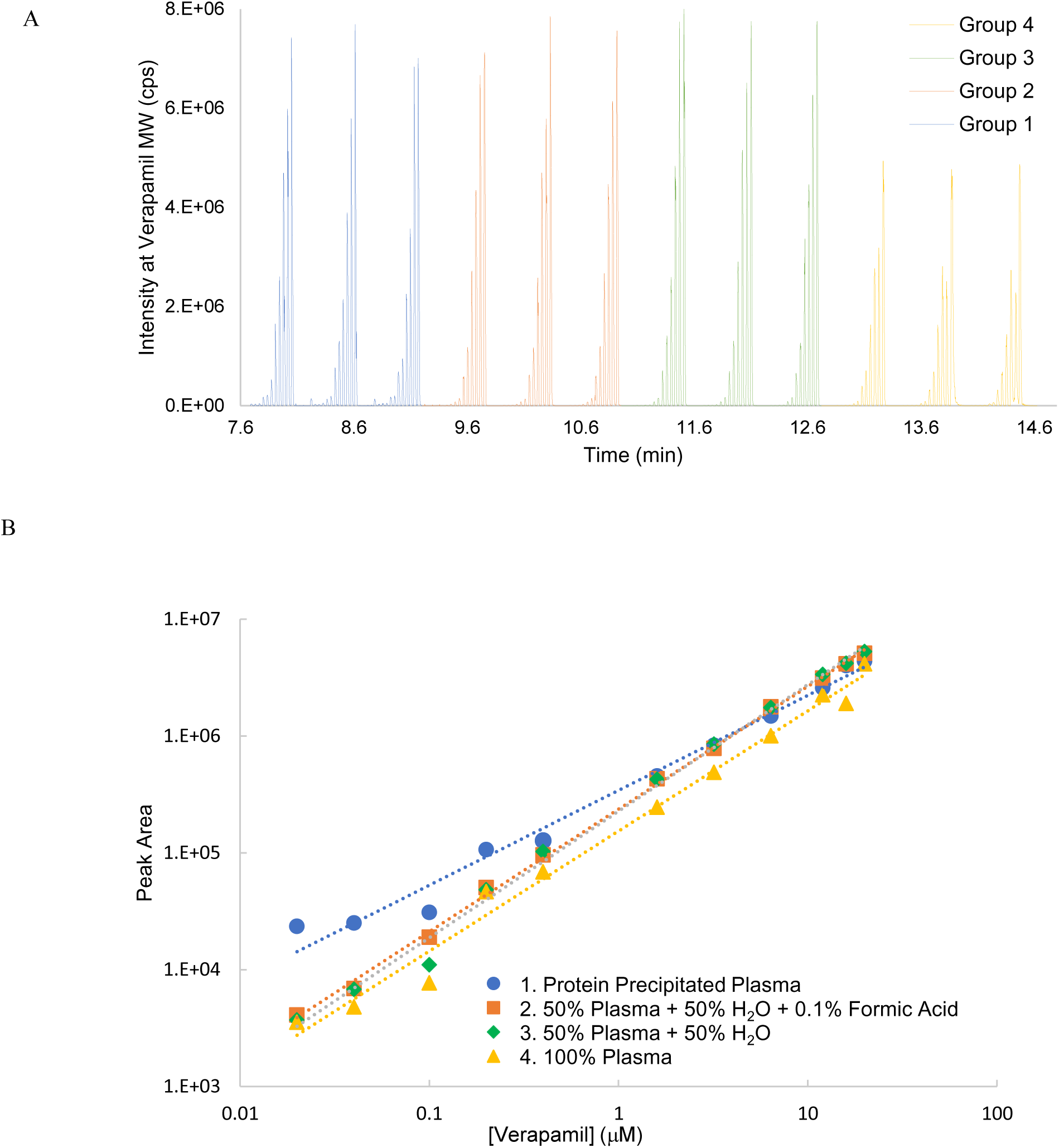
Matrix tolerance tests: standard curve samples of verapamil detected directly by ADE-OPI-MS. Four sets of verapamil standards were prepared in pooled human plasma, with final concentrations of each sample in the range 20 nM – 20 µM. Group 1 samples were subjected to conventional protein precipitation and centrifugation steps; Group 2 samples were mixed with equal volumes of water containing 0.1% formic acid; Group 3 samples were mixed with equal volumes of water; Group 4 samples were in the neat plasma (e.g., untreated). All samples were injected in triplicate. (A) Raw MS traces; (B) standard curves derived from each of the four conditions.

The assessment of enzyme reaction kinetics can be critical in the understanding of biological and chemical processes. For example, pharmacological assay kinetic parameters (e.g., enzyme/substrate concentration, incubation time, etc.) must be determined as a prerequisite to creating robust assays. Conventional approaches for such assays employ photometric technologies, specifically fluorescence plate readers. These provide high throughput, but are limited by dynamic range (usually less than 2 orders of magnitude), and detection often requires the use of special probes that may introduce experimental artifacts. MS detection affords a label-free and universal method with high sensitivity and wide dynamic range (typically 3-4 orders of magnitude). Conventional LC-MS, however, is limited by analysis time and sample processing steps that make it impractical for intensive, real-time kinetic studies. The performance attributes of ADE-OPI-MS provide the potential to measure real-time kinetics in situ, overcoming the challenges of photometric and LC-MS approaches, while allowing sampling at photometric reader speed, a wide dynamic range (3-4 orders of magnitude), tolerance for complex sample matrices free of sample clean-up, and low sample consumption. To examine a kinetic application, we conducted a biochemical experiment to measure the in situ kinetic metabolism of the CYP2D6 substrate dextromethorphan (DXM) by human liver microsomes (HLM). Both the depletion of the substrate (DXM) and the formation of the metabolite dextrorphan (DXO) were monitored over a 30-minute reaction (Fig. 3A). At each sampling event, a single 2.5 nL droplet was ejected from the 40 µL microplate well reaction volume, leaving the bulk reaction volume essentially unaltered. Repeated sampling from the same well eliminates well-to-well variability and enhances reproducibility as compared to conventional methods, wherein multiple replicates are required to average out data variability. Notably, the crude reaction samples (containing high concentrations of both protein and salt) were directly injected into the ADE-OPI-MS platform without sample alteration or cleanup steps. Peak areas of DXM for all the wells over the full time period afforded linear plots for the 6 different concentrations (Fig. 3B), demonstrating that the ADE-OPI-MS technology provides both quantitative and highly precise data. Similarly, DXO formation increased linearly with time for each of the substrate concentrations (Fig. 3C). The derived reaction rates were subsequently fitted to a Michaelis-Menten model to determine the K_m_ for metabolism of DXM to DXO CYP2D6 in HLM (Fig. 3D). At 3.79 µM, this K_m_ is within the reported range of 2.5-15.2 µM for the CYP2D6-mediated metabolism of DXM (*23*). These high-quality results, encompassing 456 data points (quantitative measurement of both substrate depletion and product formation; vastly over-sampled), were obtained in 28 minutes by one analyst.

**Fig. 3.**
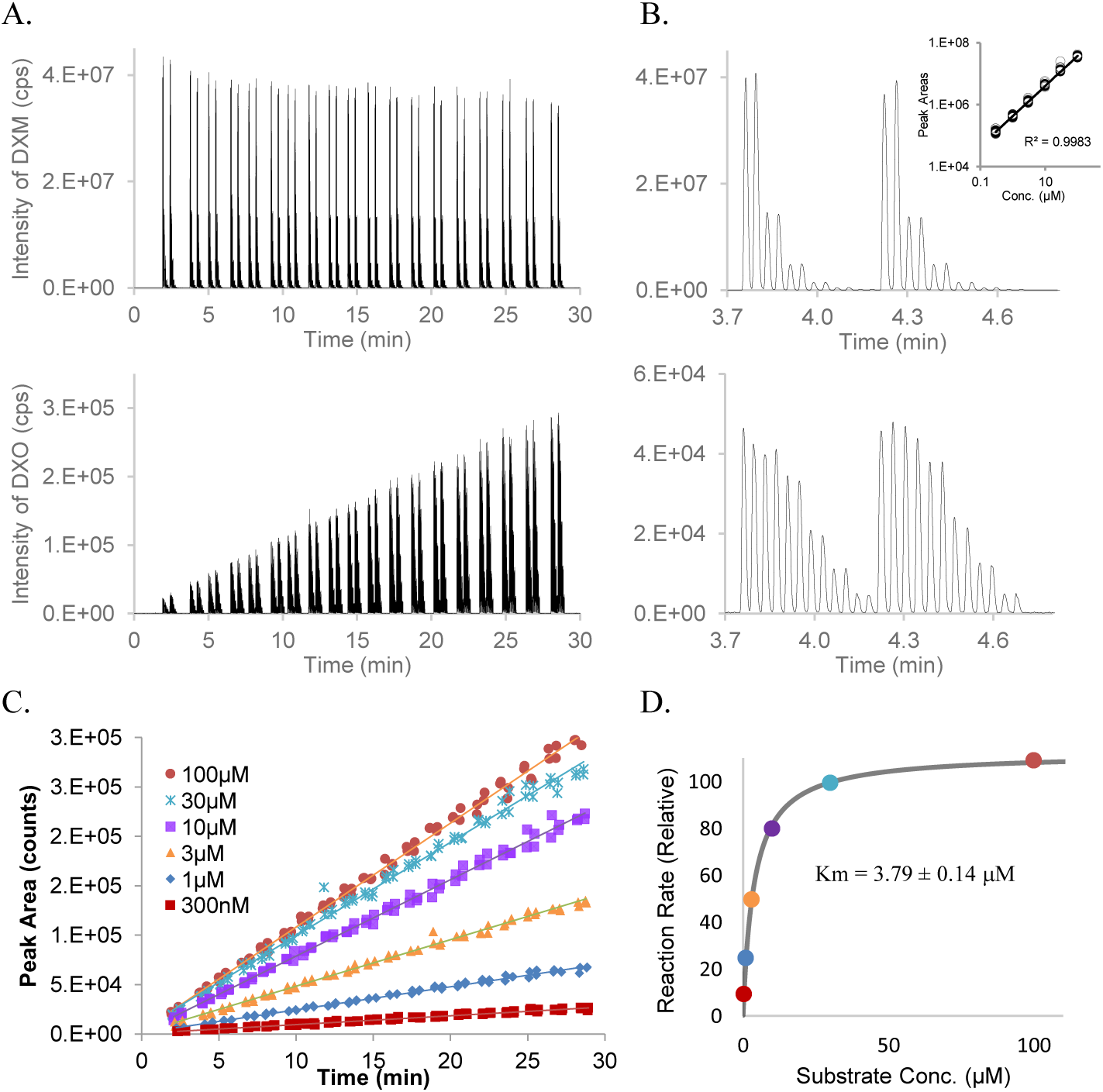
A demonstration of in-situ kinetics monitoring with the ADE-OPI-MS platform. (A) ADE-OPI-MS time traces for a CYP2D6 enzymatic reaction showing the remaining substrate dextromethorphan (DXM) and the O-demethylation metabolite dextrorphan (DXO). Each pattern reflects 6 dextromethorphan concentrations, run in duplicate. (B) Expanded view showing two sets of the 12 DXM substrate peaks (top) and the corresponding DXO product peaks (bottom) at initial reaction stage (∼4 min into the reaction, with small turnover of substrate DXM due to the low reactivity at room temperature). (C) A plot of the peak areas for formation of metabolite product (DXO) for the range of DXM substrate concentrations, measured over a 30 min reaction time. All six plots returned R^2^>0.99. (D) A Michaelis-Menten fit of the formation rate. The Km for metabolism of DXM to DXO by CYP2D6 in HLM is shown.

The ADE-OPI-MS platform is well suited for sample-intensive pharmacology applications. Acyltransferase was used as a model enzyme to demonstrate the potential utility of this platform to support typical high-throughput screening (HTS) pharmacology assays. Fig. 4 presents an example of an acyltransferase activity assay with a kinetic readout; reaction rates were determined from this set of samples via ADE-OPI-MS and compared with a conventional plate reader-based method employing derivatization with commercial CoA Green kit (Fig. 4G). For assays using high concentrations (>4 µM) of acyl-CoA substrate, similar inhibition profiles were observed with both methodologies. However, at lower concentrations of substrate (0.1-4 µM) ADE-OPI-MS was able to accurately measure depletion of substrate (see the region below 1 µM), while the plate reader returned a false formation rate due to the extra time (15 minutes according to the assay protocol) needed for coupling of the CoA reagent to enable detection (Fig. 4G). These results demonstrate that conventional plate reader-based approaches requiring coupled fluorescence probes are prone to artifacts and may not be able to determine true enzyme kinetics due to the confounding kinetics of the coupling reaction. The ADE-OPI-MS platform provides the benefits of label-free MS detection while matching plate reader sampling speed. Significantly, the ADE-OPI-MS platform can detect multiple analytes or chemical entities simultaneously, as demonstrated here for both substrate and product analyses. This is not feasible with conventional plate readers, for which specific labeling reagents need to be developed, and multiplexing may be limited by the specificity and potential interferences between labels. The application of this technology to another pharmacology HTS assay (choline transporter activity assay, Fig. S8 and Table S2), as well as ADME assays (CYP drug-drug inhibition assay, Fig. S7 and Table S1), are described in the supplementary information.

**Fig. 4.**
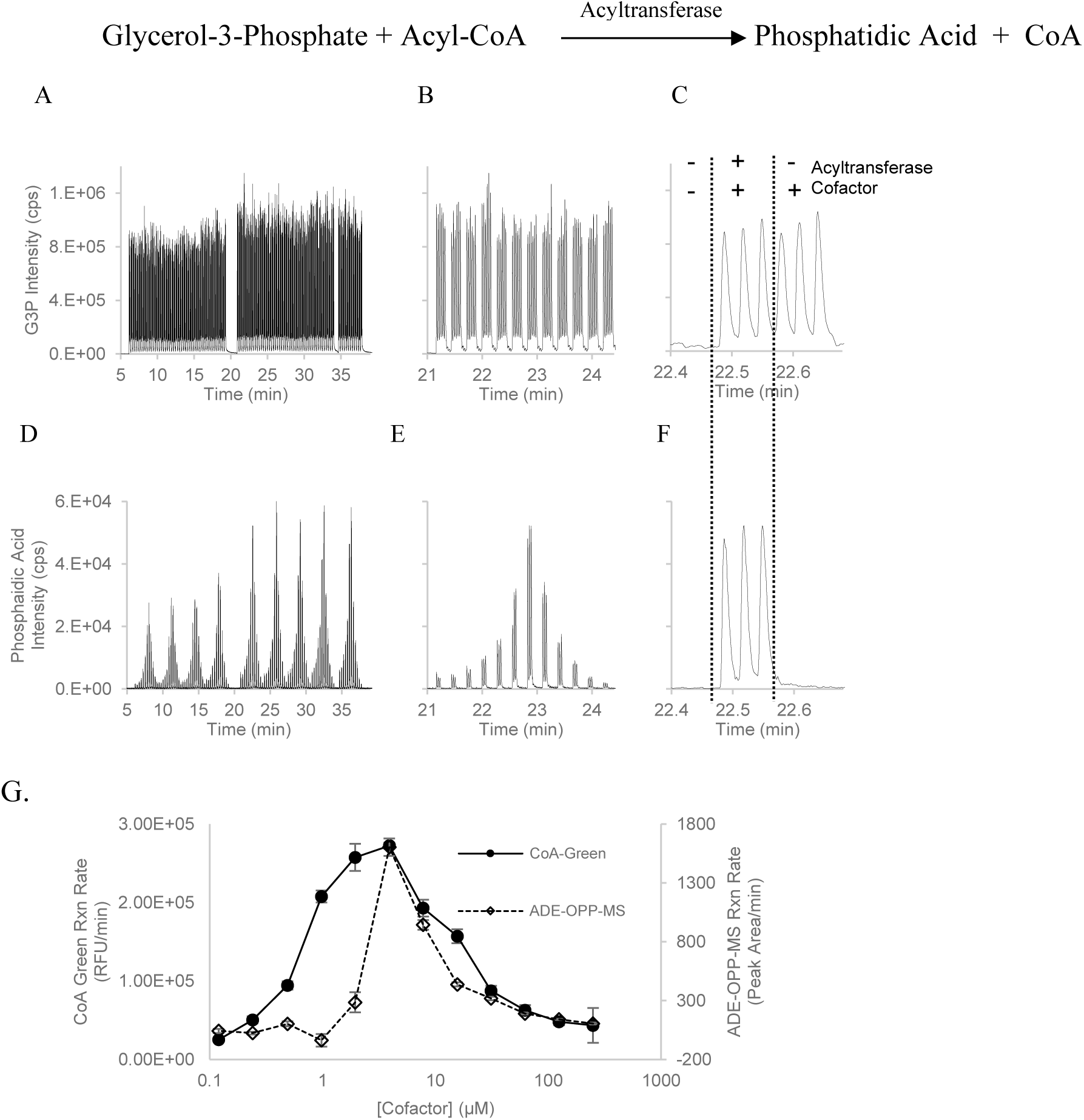
Kinetic readout of acyltransferase activity assay. 108 samples were prepared in 5 nM acyltransferase enzyme, 5 mM glyceraldehyde 3-phosphate (G3P) substrate and various concentrations of cofactor (acyl-CoA). Formation of phosphatidic acid product from the 108 samples was monitored repeatedly, 9 times within 36 minutes (D), as well as the concentration of the G3P substrate (A). Zoom-in views (B and E) of one of the nine readings, show the 12 groups of samples, each with a different acyl-CoA concentration (ranging from 250 µM to 0.1 µM with 2X step dilution). Further zoom-in views (C and F) of 9 samples run in triplicate with the same 4 µM acyl-CoA concentration, but with and without enzyme/cofactor (see label in insert). G. Comparison of the reaction rates derived from fluorescence (BioVision Co-A #K367-100) and ADE-OPI-MS (phosphatidic acid) assays.

The ADE-OPI-MS platform is compatible with any type of mass spectrometer that is used in conventional LC-MS. In addition to the unit-resolution triple-quadruple instrument optimized for targeted quantitation applications shown above, we have also integrated a high-resolution (HR) time-of-flight (TOF) instrument into an ADE-OPI-MS system to create an HR variant (ADE-OPI-HRMS). When operating in full mass profiling mode, ADE-OPI-HRMS becomes a label-free, high-throughput sampling, high-content reader. To gauge ADE-OPI-HR-MS utility, we analyzed samples from plate-based miniaturized parallel medicinal chemistry synthesis. With simple post-acquisition data mining, the formation of the targeted product versus remaining starting material can be monitored simultaneously. A chemical reaction optimization campaign is shown in Fig. 5, where multiple experimental conditions were assessed for a specific coupling reaction. The set of 96 reactions was evaluated by both conventional LC-MS and ADE-OPI-MS for the same analytes. Notably, the ADE-OPI-MS platform generated the analytical results 60-fold more rapidly (5 minutes versus 5 hours) and utilized 1000-fold less sample (sample injection volume 2.5 nL versus 2.5 µL). For synthesis efforts in drug discovery, this translates to significant cost and time savings. Further, the ESI source enables broad coverage of chemical space (*24*).

**Fig. 5.**
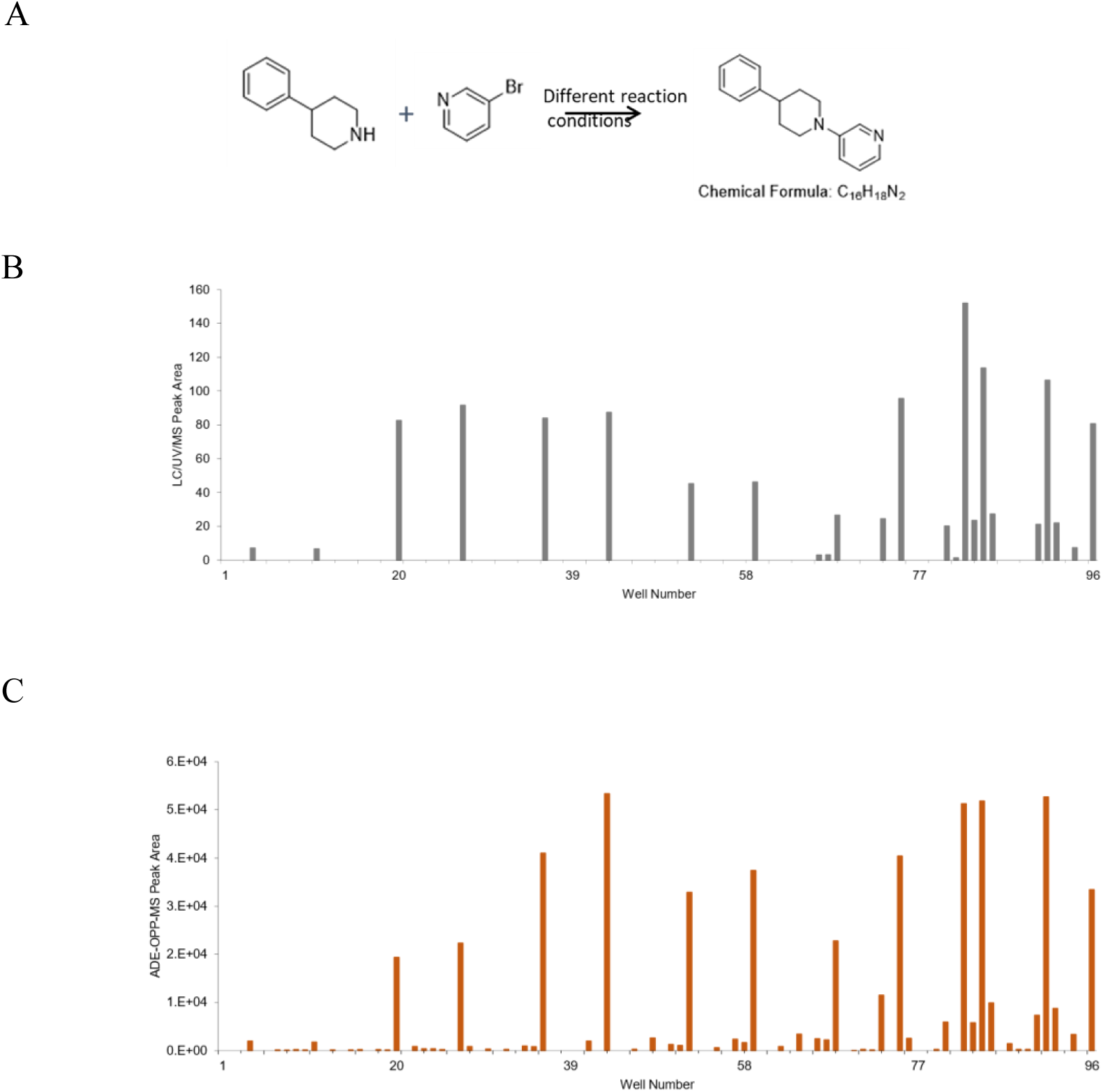
(A) Chemical reaction pathway for a simple coupling reaction with a targeted product C_16_H_18_N_2_ (*25*). (B) Mass detection of product formation with 96 different reaction conditions using LC-UV-MS. Peak areas of the target product were plotted against the well numbers. Each injection consumed 10 μL of the reaction and each sample took 3 minutes for analysis. Overall, it required 5 hours to analyze all 96 reactions using conventional LC-UV-MS. (C) The same set of samples were analyzed using ADE-OPI-MS. Only 5 nL of the sample was consumed for each reaction. The 96 samples were injected continuously, and total analysis time was 5 minutes. Shown here is the extracted ion chromatogram of protonated product (*m/z* 239.15).

This initial set of results demonstrates that the ADE-OPI-MS technology, via its performance characteristics and ease of use, has the potential to greatly expand fundamental mass measurement as the analytical technique of choice for experiments where accurate and sensitive analysis is critical. As outlined in Table 1, ADE-OPI-MS offers high speed, high throughput, and miniaturized experimentation; it will therefore allow broader adoption and utility of MS technology, as well as enhancing the experimental quality of the results. The sheer speed of analysis and applicability to a wide breadth of analyte classes allows a given instrument to perform millions of determinations over the course of a few months, in support of a wide array of sample types. The approach’s robustness and its ability to analyze unprocessed samples with minimal methods development greatly enhance the speed of the entire experimental workflow, while facilitating miniaturization and preservation of original samples. In conclusion, the ADE-OPI-MS technology allows for dramatic performance increases for current LC-MS applications, and opens the door for utilizing MS in new application space.

**Table 1.**
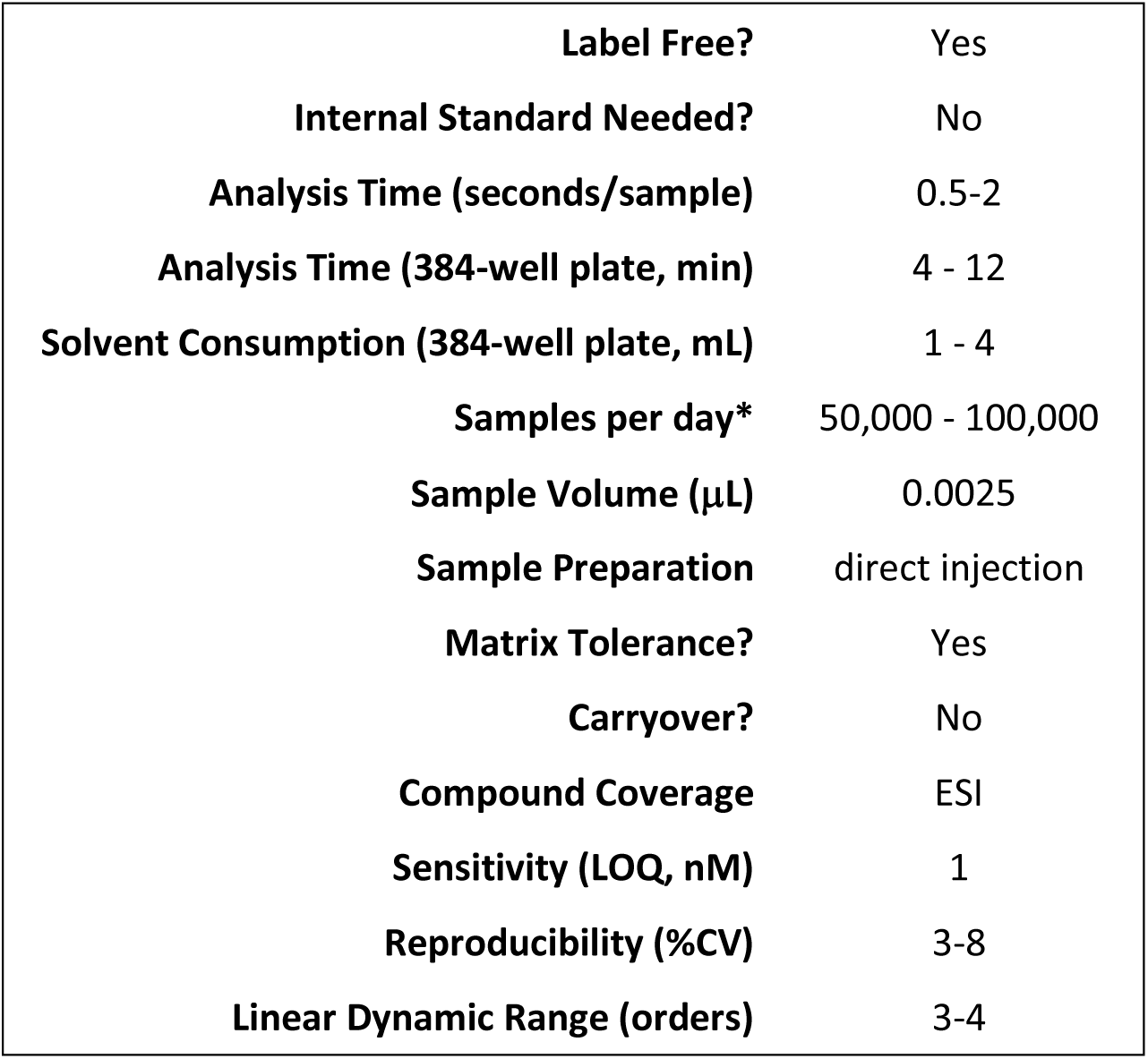
Overview of characteristics for ADE-OPI-MS. *extrapolated from 384-well plate analysis time.

## Supporting information

Movie S1: ADE-OPI-MS system in operation

## Acknowledgments

We thank Dr. Mark Noe for support in helping establish the collaboration between Pfizer, Sciex, and Labcyte to develop this technology at Pfizer’s Groton, CT site. We thank E. J. Corey for his guidance and insight and Drs. D. Cox, L. Burton, K. Brighty and M. Blauwkamp for helpful discussions and reviewing the manuscript. We thank Mike West for sharing LC/MS data used in Sup.Fig.7 for comparison.

## Author contributions

H.Z., C.L., W.H., J.L. designed the studies and experiments. L.G, S.S.D. provided the ADE setup. C.L., T.R.C., D.W.A. provided the OPI setup. H.Z., C.L., W.H., J.L., L.G., L.A., S.N., K.D., L.F.L., C.M.S., D.W.A., T.R.C., S.S.D. acquired the data. H.Z., C.L. performed data analysis and interpretation. H.Z., C.L., L.G., S.S.D., M.D.T. wrote and revised the manuscript. H.Z., C.L., S.S.D., M.D.T. supervised the study.

## Supplementary Materials, for

### Supplemental Text

#### Instrumentation, Materials and Methods

##### ADE-OPI-MS Instrumentation

The following section will discuss salient features of the components and complete ADE-OPI-MS system.

###### The ADE Technology

Acoustic droplet ejection (ADE) technology is optimized to deliver small volumes of solution from source to destination using acoustic energy. Historically, this technology has been integrated with an inverted microplate held on a destination stage for drug discovery applications (compound reformatting, dose response setup) and for screening applications (pharmacology, ADME), proteomics and genomics (*26*). At each well of the source microplate, a piezoelectric transducer with an input waveform at 10 MHz center frequency (CF) is used to generate a focused ultrasonic pulse which propagates through the coupling fluid (figure not shown) into the walls and contents (Fig. 1A), (*27*). This ultrasonic pulse is reflected at the interfaces (including the fluid meniscus) and returns to the piezoelectric transducer for real-time processing to audit the microplate (e.g. to measure the: fluid meniscus position; the fluid meniscus tilt; the fluid impedance). At greater burst amplitude acoustic energy is focused at the sample fluid surface and acoustic pressure is applied to form a mound at the fluid meniscus. These ultrasonic pulses reflected by the fluid menisci are processed by dynamic fluid analysis (DFA) algorithms to determine droplet ejection parameters.

Next, a burst pulse is then applied to acoustically transfer droplets. Droplet diameter is in the range 120 – 360 μm (1 - 25 nL in volume), with a typical droplet velocity of 1 m/s that is directed to the target (*28*). The ADE repetition rate of this process is 200-800 Hz, transferring a droplet train from the same sample well at an effective infusion flow rate of 30 – 75 μL/min (Figure S1A and S1B). The typical injection volume is a single droplet (2.5 nL), and a droplet train of 10 or more droplets is possible. ADE allows for precise and consistent transfer of a selected number of sample droplets into the OPI for analysis as a single peak. The entire system operates at room temperature and pressure, it can be used with a variety of solutions with equivalent accuracy and precision (Figure S2).

###### Materials

Sodium hydroxide (J.T. Baker, 5674-02)

Echo® qualified 384-well polypropylene microplates (Labcyte Inc., P-05525)

384-well clear-bottom polystyrene microplates (Greiner Bio One, 781096)

Synergy H4 Hybrid multi-mode microplate reader (BioTek Instruments, Inc.)

All other chemicals and reagents were purchased from Sigma-Aldrich.

###### Method

In the droplet volume verification method, each test solution was prepared with 0.15 mM sodium fluorescein as a fluorescence tracer dye. The ADE liquid handler was setup for twenty droplet (50 nL) transfers from each test solution prepared in an Echo qualified 384-well source microplate into a clear-bottom 384-well destination microplate. The 384-well source plate for each solution was prepared in a quadrant fill pattern, with 96 wells filled to each of four volumes: 15, 20, 30, and 65 µL. Following the transfers, the clear-bottom microplates were back-filled with 50 µL per well of 10 mM sodium hydroxide using a conventional bulk filler, centrifuged for 1 minute at 1,000 rpm and then incubated for 30 minutes at room temperature. Next, the microplate was read on the Synergy fluorescence reader to determine the fluorescence level in each well. The fluorescence level was compared to a standard curve to determine the transfer volume for each well. Fig. 2 shows the average transfer volume and coefficient of variation (CV) results for the following fluids: glycerol (0-60%, 10% steps), dimethyl sulfoxide (DMSO) (70-100%, 5% steps), fetal calf serum (FCS) (0-100%, 20% steps). Triton X-100 (0-200% CMC, 0, 5, 14, 200%). DMSO dilutions are in Milli-Q H_2_O, the remaining dilutions were prepared with 1X phosphate buffered saline (PBS). Triton X-100 dilutions are 0.001%, 0.003%, and 0.042% (v/v), these concentrations represent 5%, 14%, and 200% of the critical micelle concentration (CMC), respectively. Fluid calibrations for organics including acetonitrile (0-100% in H_2_O) and methanol (up to 50% in H_2_O) are also available (data not shown).

###### The Open Port Interface (OPI)

The OPI sampling interface uses a vertically aligned, co-axial tube arrangement enabling solvent delivery through a tubing annulus to a capture region, the open-port, Fig. S1C, (*17*). The inner diameter (ID) of the outer tube (stainless steel, electrically grounded) is 950 μm, and the inner capillary (outside diameter: 800 μm, and ID: 250 μm) is recessed by 0.3 mm within outer tube to define a cylindrical fluid vortex volume. In this study, an OPI (open port facing down) is positioned upside-down above the source well of a microplate (1-3 mm clearance) and aligned to the ADE transducer axis to capture acoustically dispensed sample droplets. A gear pump delivers a typical liquid chromatography (LC) mobile phase with flow rate in the range 300-600 μL/min to the open port.

The inner capillary (L = 30-60 cm, PEEK or PEEKSil) connects from the open port to the ESI electrode of a Turbo V ion source (200 μm ID, stainless steel) of the MS (SCIEX Triple Quad^TM^ 6500+ System or SCIEX TripleTOF^®^ 6600), with ion spray voltage at 5500 V and 300°C source temperature. Expanding gas at the nebulizer exit (ESI source nebulizer gas, fixed at 90 psi, Fig. S1) draws a fluid stream by the Venturi pressure drop created at the ESI nozzle (630 μm ID).

The ESI electrode protrusion from the nozzle is 300 μm. The OPI carrier flow rate is optimized to balance the nebulizer flow rate to achieve a stable vortex in the droplet capture region.

Samples enter the capture region, mix and dilute in the vortex and flow within the capillary to the ESI electrode and nozzle for detection by the MS.

###### The ADE-OPI-MS system

The ADE-OPI-MS platform shown in Figure S1 consists of an externalized transducer assembly from an Echo 555 acoustic liquid handler (Labcyte, Inc., San Jose, CA), a breadboard XY stage with a source microplate gripper (Labcyte, Inc., custom), an open-port probe sampling interface connected to both a carrier solvent pump and a transfer capillary leading to the standard IonDrive^TM^ Turbo V ESI source of a Triple Quad^TM^ 6500+ System or TripleTOF^®^ 6600 (SCIEX).; Fig.S1 and the Supplemental Movie S1). An XY stage carrying the source plate provides for rapid translation between source wells enabling high throughput sample processing at greater than 3 samples per second. This ADE breadboard system was operated with customized Cherry Pick client and server software (v.2.5.MS, Labcyte Inc.). The mass spectrometer was operated with standard software (SCIEX Analyst 1.6.2). Movie S1 shows the unit in operation. Another breadboard version of the ADE-OPI-MS platform was used for some part of this study, consisting of a standard ATS Gen 4+ system (EDC Biosystems, Fremont, CA). For this system, the rotation function of the target-plate gripper on the ATS system was disabled. A 3-D printed OPI-holder with external dimensions of a standard microplate was loaded in the target-plate gripper to position the open port capture region with a gap of 2 mm above the source microplate. The x-y position of the OPI was set in software to align the OPI capture region to the acoustic transducer axis. In a separate step, the source well is also aligned to the acoustic transducer axis. The ATS-100 software was used to control acoustic ejection.

##### General approaches utilized to generate analytical samples, analysis and data processing used for data presented through this work

###### Preparation of Plates for ADE-OPI-MS Analysis

Prior to loading in the system, source microplates with sample fluid were centrifuged (2100 RCF for 5 min) to remove gas bubbles and to provide a consistent fluid meniscus shape. Next the source microplates were deionized to minimize droplet deflection by electrostatic charge within the source well. The full range of ADE compatible labware includes a 6-well reagent reservoir, acoustic tubes and microplates with 96-, 384-, and 1536-well formats. For each source plate there was a defined operating fluid volume range (e.g. 20 – 65 μL for 384-well).

###### Sample Analysis with ADE-OPI-MS

With the OPI device aligned to the acoustic transducer axis in x-y (+/− 100 μm) and the vertical position of the OPI set 1-3 mm above the top surface of the source plate, the ADE droplet sampling protocol could begin. The OPI carrier solvent flowrate was optimized at the fixed nebulizer gas setting (i.e. 90 psi) to achieve the optimal stable vortex (best reproducibility and sensitivity). The delay time between samplings was adjusted according to the MS peak-width to maintain the baseline separation between adjacent signal peaks. The sampling protocol steps included: loading the sample plate into the gripper of the x-y stage, stage translation to position a selected source well above the acoustic transducer, vertical positioning of the transducer to focus at the sample fluid surface, dynamic fluid analysis (DFA) to determine acoustic ejection parameters and final droplet transfer from the source well sample fluid into the capture region of the OPI. The sample ejection volume is adjusted by changing the number of droplets sampled at the selected dispensing repetition rate (20-800 Hz). Multiple ejected droplets diffuse, mix and merge in the capture region vortex to increase ejection volume.

###### Data Processing

Data was post-processed by parsing the MS raw data file (.wiff) and the acoustic ejection log to correlate droplet transfer events with ion count peaks. The delay time from a droplet ejection event to the detection on an ion count peak is 3 – 10 seconds, depending on the carrier solvent flow rate (200 – 600 μL/min) and length of the transfer capillary (30 – 60 cm). Post-processing software generates an output file tagging each ion count peak with a source well location. Data processing can be completed offline and takes around 2 min per 384-well plate using a laboratory PC.

##### ADE-OPI-MS Analytical Performance

First, we have explored the potential to increase overall sample analysis speed by deploying multiplex analysis. Figure S3 shows a trace of 384 samples recorded in < 170 seconds (2.2 Hz) by multiplexing four different compounds (omeprazole, quinidine, midazolam, bupropion) ejected sequentially from individual wells. Importantly for this multiplexed approach, the precision remained high with CV% between 3% to 8% for these model compounds.

Analytical performance of the ADE-OPI-MS was gauged using the typical analytes, propranolol and midazolam. Sensitivity was measured with a droplet ladder (1 to 10 droplets, 2.5 nL to 25 nL) of 25 nM propranolol in aqueous solution (Figure S4A). The calculated limit of detection (LOD) for propranolol was 1 attomole loading based on the last peak with 62.5 attomoles (2.5 nL x 25 nM, as highlighted in Fig. S4A). The sensitivity and linear dynamic range (LDR) were further evaluated by ejecting a single sample droplet (2.5 nL) across standard curves ranging from 10 nM to 10 mM of propranolol (Figure S4B) and midazolam (Figure S4C). These calibration curves shown indicate the LDR is at least 3 orders of magnitude without use of internal standard.

Next, we explored the possibility of the direct loading of samples containing detergents with ADE-OPI-MS. Due to the low sample loading amount (<10 nL) and the high flow rate of carrier solvent, the sample matrix is significantly diluted (∼1000x) within the capture region. Figure S5A shows a raw MS traces from two alternating 17-aa peptides (New England Peptide Inc., Gardner, MA), shown in pink and blue respectively, at three different concentration levels: 500 nM, 250 nM, and 125 nM in 50 mM HEPES buffer (Lonza, Walkersville, MD) with and without a commonly used detergent, 0.01% Tween 20. In Figure S5B, a histogram shows peak areas for the model peptides. Thus, we do not observe significant ion suppression with the addition of Tween 20 detergent. The process of sample cleanup is a major bottleneck in high-throughput bioanalysis because it is compound dependent and may vary with different matrices. This “online dilution” feature allows us to eliminate or significantly simplify the sample preparation.

After obtaining highly accurate and precise data across a wide dynamic range for small molecules and peptides, we investigated larger molecules. In Figure S6, we show the analysis of an antibody standard (MW∼150K, Waters, Milford, MA) with alternating five droplet (12.5 nL) injections of the standards at two concentrations, 667 nM and 67 nM in aqueous solution. The estimated LOD is <1 fmole loading.

##### Further ADE-OPI-MS Applications Supplemental Text with Materials and Methods

We tested the ADE-OPI-MS platform for a key drug properties application, drug-drug interaction (DDI) profiling (Fig. S7). One of the first and most important DDI studies involve profiling cytochrome P450 (CYP) DDI potential. CYP DDI assays monitor the formation of metabolites with co-dosing of test compounds where a decrease in the formation of the metabolite indicates inhibition of the corresponding enzyme and that there is a potential risk of perpetrator DDI when co-dosed with other drugs metabolized by that enzyme. These are of great importance and concern due to the severity of DDI’s related to altering CYP activity (*29–30*). In this study, 16 compounds were tested for potential inhibition of 3 CYP enzymes, evaluated in tandem using the specific substrates: midazolam, dextromethorphan, and tacrine for CYP3A4, CYP2D6 and CYP1A2, respectively.

###### Standard experiment conditions

Three specific substrates midazolam (for CYP3A4), dextromethorphan (for CYP2D6), and tacrine (for CYP1A2) were mixed with pooled human liver microsomes (HLM, 0.1 mg/mL) at final protein concentration of 2, 5, 2 µM respectively. Test compounds were spiked in the samples to achieve a final concentration of 0, 30 nM, 100 nM, 300 nM, 1 µM, 3 µM, 10 µM, 30 µM each. The reactions were initiated by the addition of reduced nicotinamide adenine dinucleotide (NADPH) at 1.2mM) to the HLM samples. The reaction was quenched after 8 minutes with addition of each volume of acetonitrile. The samples were next centrifuged and then injected on both LC-MS and ADE-OPI-MS platforms and the corresponding metabolites (OH-midazolam, dextrorphan, and OH-tacrine) which were monitored simultaneously over a range of test compound concentration (Fig. S8). The Z’ factors, IC_50_ and EC_50_ data show good concordance, but ADE-OPI-MS operates without the need for an internal standard and with greatly reduced solvent, sample and time (Table S1).

###### Choline uptake assay

We also demonstrate a first implementation of the ADE-OPI-MS platform in a high-throughput screening (HTS) drug discovery application (Fig. S8). The choline transporter (CHT) is an attractive target for neurological disorders such as Alzheimer’s and attention deficit hyperactivity disorder (ADHD) as it is known to mediate synthesis and distribution of the critical neurotransmitter acetylcholine. Conversely, cholinergic dysfunction is also associated with attention deficiencies and compromised motor neuron function. In order to screen for modulation of CHT activity by a candidate drug molecule, a cellular uptake assay was developed and validated to monitor the uptake of the deuterated (D-9 labelled) choline, as choline has high endogenous background levels in the HEK293 cell line. HEK cells expressing the high affinity CHT were used for this assay. Known modulators hemicholinium-3 (HC-3, inhibitor) and staurosporine (STS, activator) (*22*) were tested at a range of dose concentrations. Due to high background signal from endogenous choline, D-9 labelled choline was used as substrate.

In 384-well poly-D-lysine coated plates, 50,000 cells per well were plated in 50 μL media and recovered overnight. The cells were rinsed and equilibrated for 30 minutes in 50 μL HBSS buffer and then pre-incubated with test compounds for 15 minutes. A final concentration of 100 μM d9-Choline substrate was added and uptake progressed for 15 minutes. Substrate was removed by aspiration and the cells were washed two times with HBSS buffer. The wells were then extracted with 30 μL of HPLC grade acetonitrile:methanol:water (2:2:1).

###### ADE-OPI-MS experimental conditions

OPI carrier solvent flow: methanol (0.20 mL/min), sample ejection volume: 5 nL, mass spectrometer: SCIEX Triple Quad^TM^ 6500+ system. Data collection: Analyst 1.6, MultiQuant 2.1, ion source temperature: 150°C, analyte SRM Transitions Monitored (CE) D9-Choline 113.2→ 69.1 (26 eV).

###### Standard experiment conditions

Lysed cell samples were analyzed using the ADE-OPI-MS platform without further treatment or cleanup and compared to a conventional LC-MS method (*31*). Pre-incubation of HC-3 and STS showed well defined inhibition and activation profiles as expected. Both profiles generated with ADE-OPI-MS had high concordance when compared to EC50 generated via conventional LC/MS. The ADE-OPI-MS platform increased the speed 10-fold and reduced sample volume 500-fold while delivering equivalent data quality and concordance for both inhibition and activation profiles of the two modulators. Further, the ADE-OPI-MS platform results were obtained without an internal standard (IS) and with higher precision than the LC-MS methods with IS.

## Supplemental Figures and Legends

**Figure S1:**
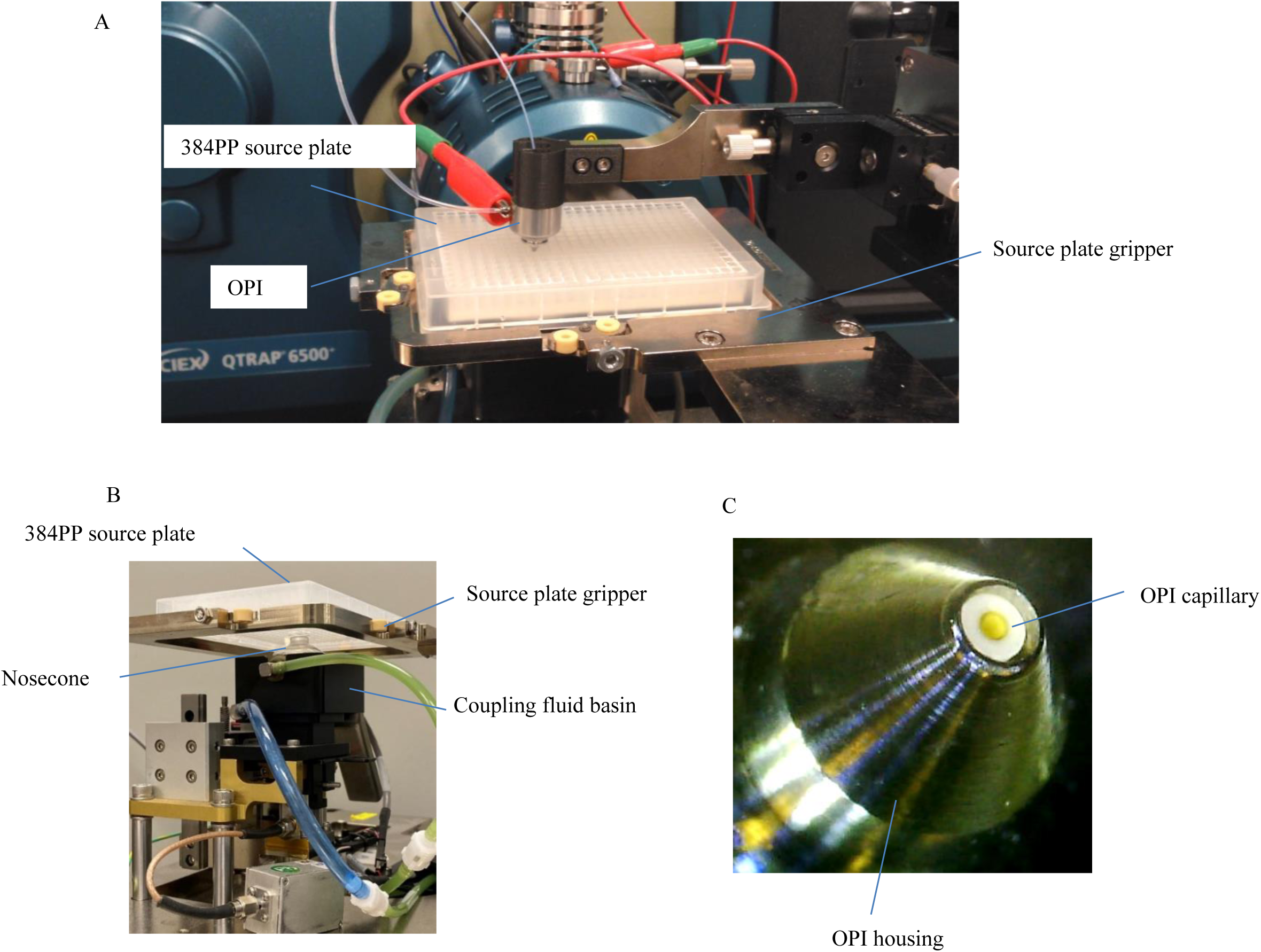
ADE-OPI-MS Breadboard System. (A) Photograph of the ADE-OPI-MS platform showcasing a 384 polypropylene source microplate in the source tray of a breadboard XY stage. The system is mechanically aligned with an Echo acoustic transducer (below stage), center of the microplate source well, and the OPI transfer capillary on a common axis. Clearance between microplate and OPI is set to 1 – 2 mm to provide for droplet placement to the center of the OPI capture region. A transfer capillary connects the OPI to a SCIEX Triple Quad^TM^ 6500+ MS with IonDrive^TM^ Turvo V ESI source. (B) Photo of ADE setup showing transducer housing, coupling fluid basin, nosecone, source plate gripper and 384PP source plate. (C) zoom-in view of OPI probe showing ID of stainless-steel outer tube and inner capillary.

**Figure S2:**
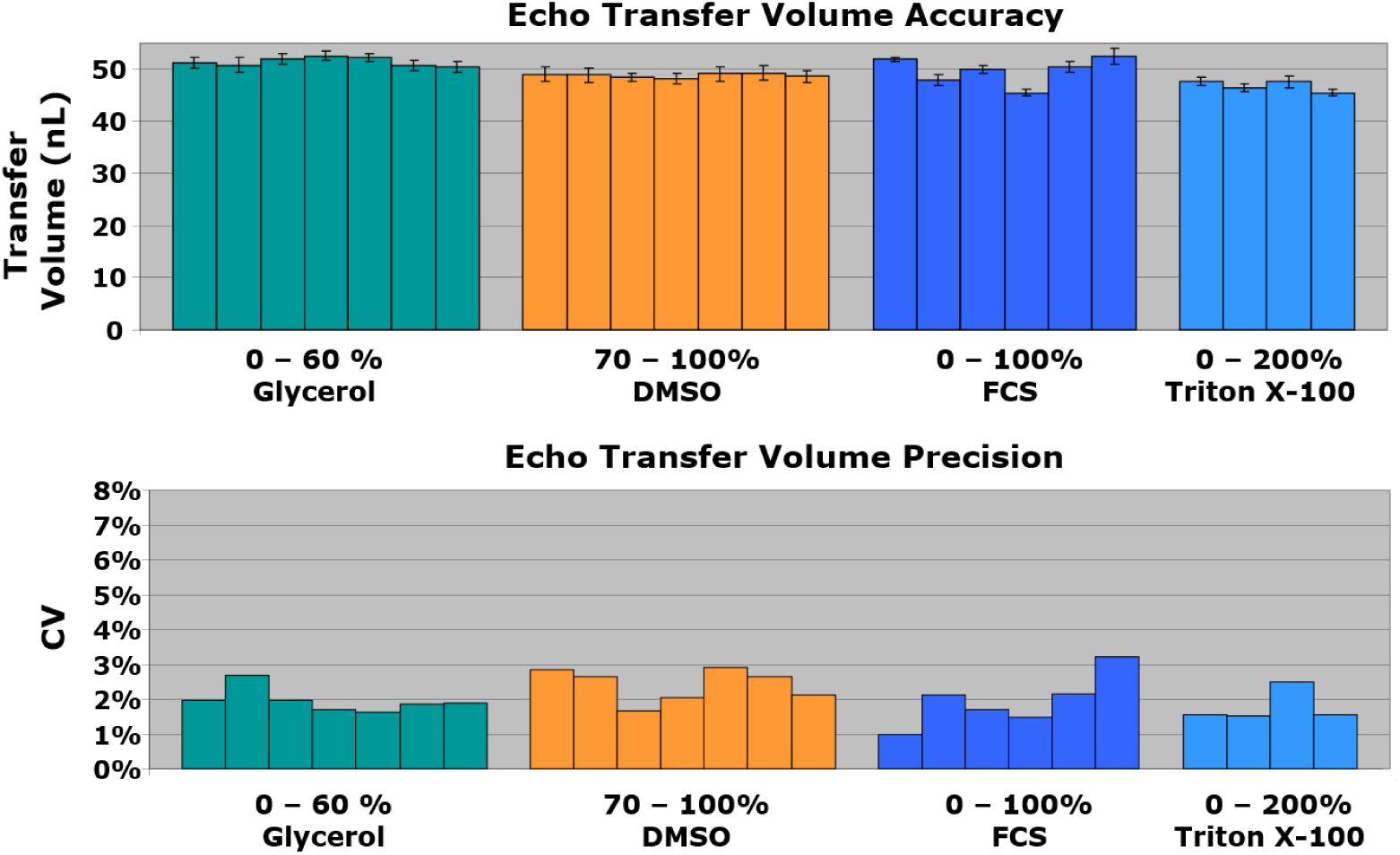
Measurement of Echo ADE Droplet Volume and Coefficient of Variation. ADE droplet transfer performance: absolute transfer volume and coefficient of variation is consistent for a wide range of sample fluids. The table shows the performance of acoustic calibrations for four fluid classes: glycerol (0-60%, 10% steps), diMethyl sulfoxide (DMSO) (70-100%, 5% steps), fetal calf serum (FCS) (0-100%, 20% steps), Triton X-100 in water (0-200% of the critical micelle concentration (CMC), 0%, 5%, 14%, 200%).

**Figure S3:**
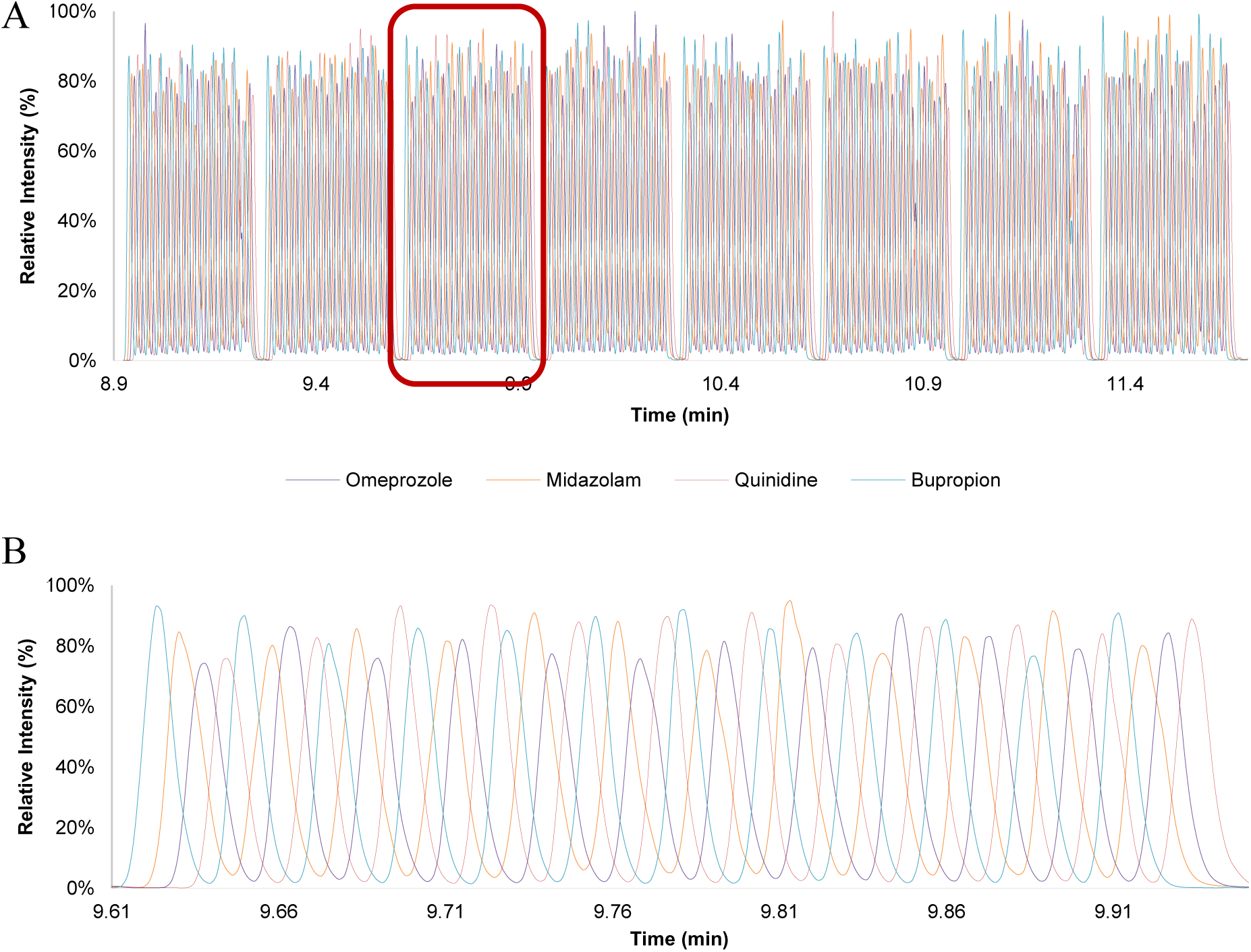
Multiplexed Analysis with ADE-OPI-MS. (A) MS trace of 384 samples measured with multiplexing in < 170 seconds (2.2 Hz) with four different compounds (omeprazole, quinidine, midazolam, bupropion) sequentially ejected from individual wells, CV in the range 3% - 8%. (B) One section of the 48 peaks shown in (A) enlarged.

**Figure S4:**
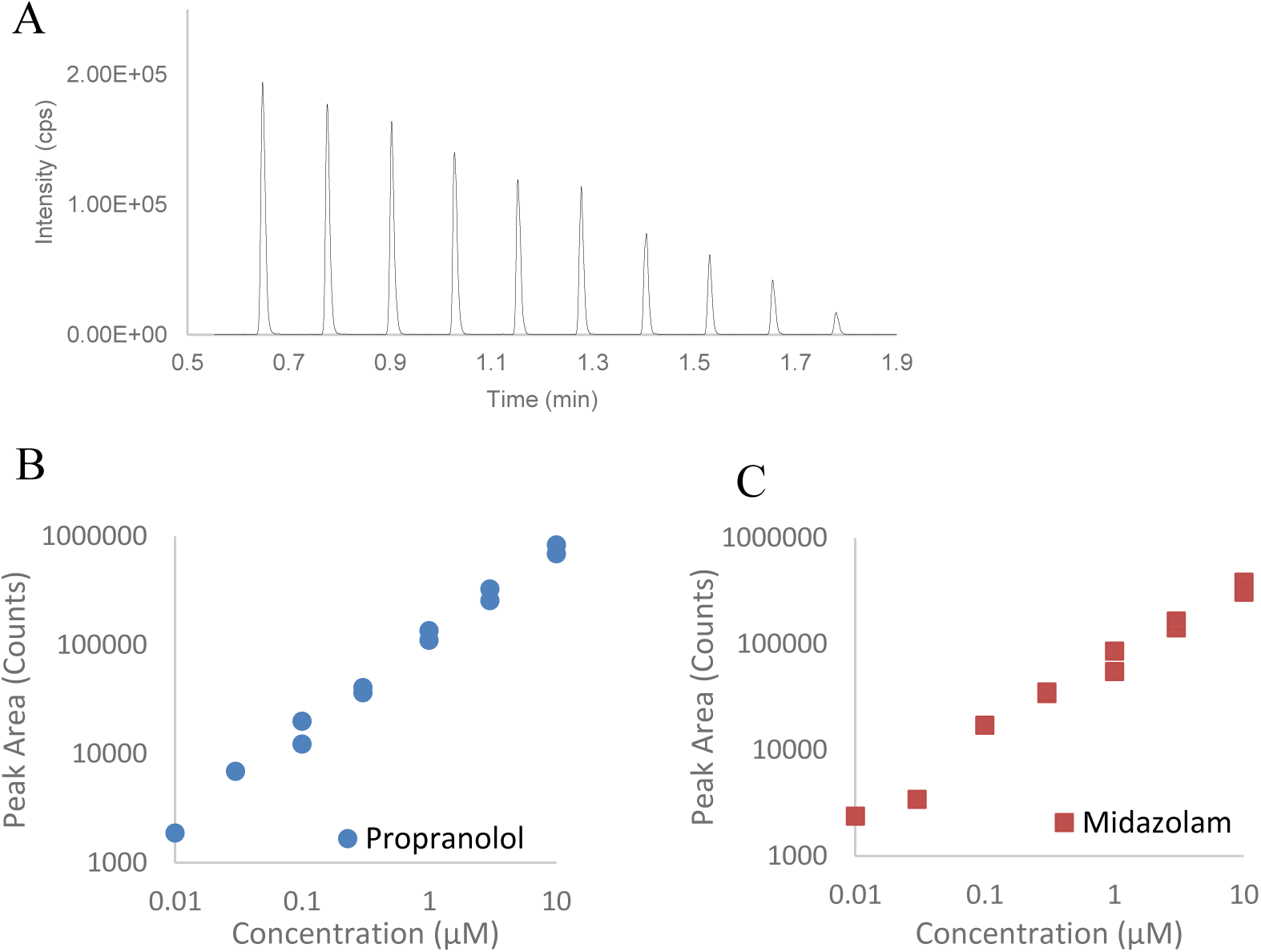
Linear dynamic range (LDR), limit of detection (LOD): (A). Analytical sensitivity was evaluated by injecting a droplet ladder (10 droplets to 1 droplet, 25 nL to 2.5 nL) of 25 nM propranolol. Sample droplets are interleaved with blank droplets, no carry-over is observed. (B) Single sample drop (2.5 nL) analysis for propranolol. (C) Single sample drop (2.5 nL) analysis for midazolam. Data in (B) and (C) performed with standard curves from 10 nM to 10 μM.

**Figure S5:**
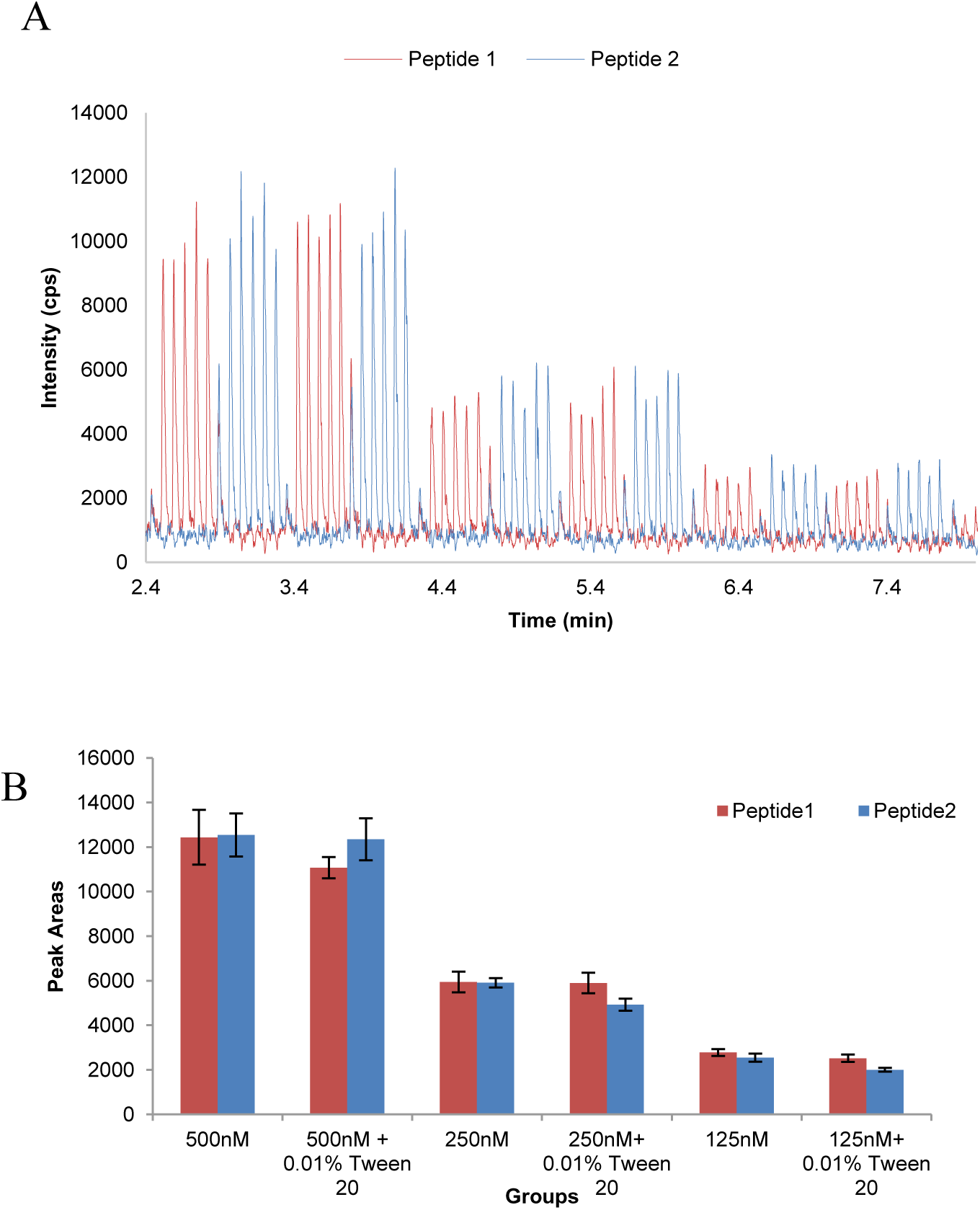
Matrix Tolerance Assessment with Detergent Samples. (A) MS traces of two alternating 17-aa peptides (shown in pink and blue respectively) at three different concentration levels: 500 nM, 250 nM, and 125 nM. (B) Histogram showing the peak areas of two model peptides in buffer alone and with a commonly used detergent, 0.01% Tween 20.

**Figure S6:**
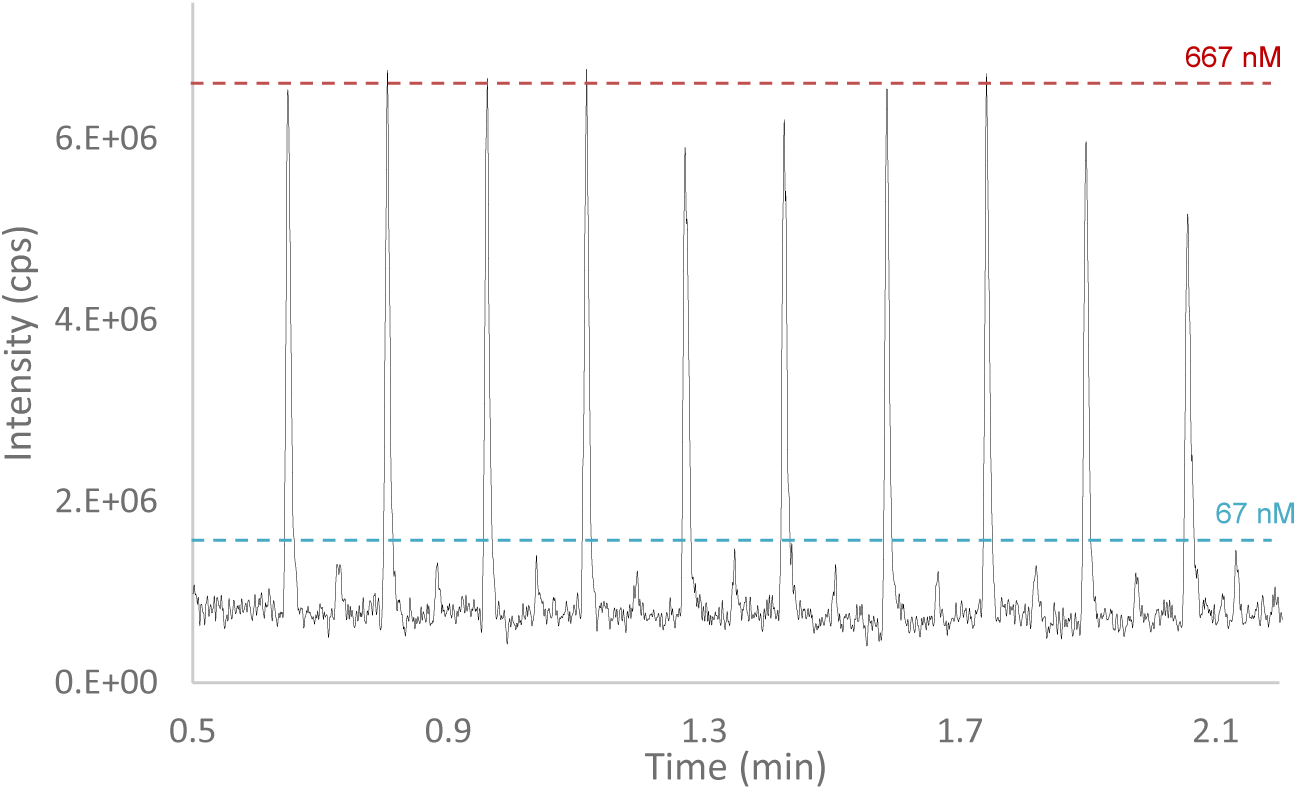
Analyte Coverage: Demonstration of Antibody Detection. ADE-OPI-MS analysis of antibody standard (MW∼150 kDa) with alternating injections of 12.5 nL standards at a concentration of 667 nM and 67 nM, respectively. The estimated LOD is <1 fmole loading.

**Figure S7:**
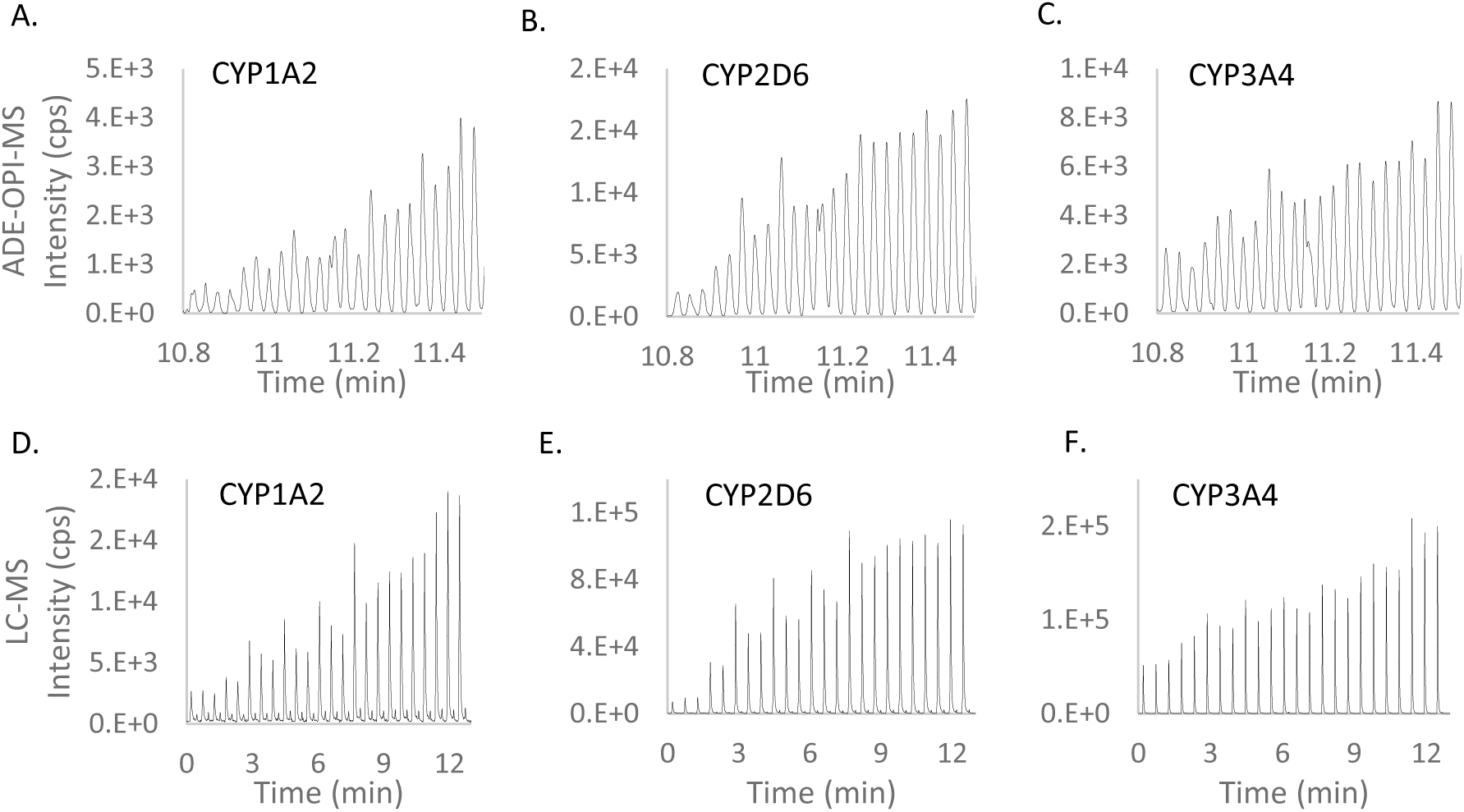
ADE-OPI-MS and LC-MS ADME DDI Experiments. MS traces of metabolite formation for one representative test compound detected by both ADE-OPI-MS (A, B, and C) and LC-MS (D, E, and F). MS traces represent the dose response of inhibition of the test compound to CYP 1A2, 2D6, and 3A4 enzymes, as shown. Test compounds were dosed at 8 different concentrations in triplicate.

**Table S1:** ADME CYP DDI IC_50_ Values (μM) Obtained using ADE-OPI-MS and LC-MS. Calculated IC_50_’s of 16 test compounds to three different CYP isoforms (1A2, 2D6, 3A4), with both ADE-OPI-MS and LC-MS platforms.

| Compound | ADE-OPI-MS |  |  | LC/MS |  |  |
| --- | --- | --- | --- | --- | --- | --- |
|  | 1A2 | 2D6 | 3A4 | 1A2 | 2D6 | 3A4 |
| A | 22.4 | 0.03 | ≥30.0 | 13.5 | 0.03 | 9.2 |
| B | ≥30.0 | 0.25 | 0.69 | ≥30.0 | 0.17 | 0.13 |
| C | ≥30.0 | ≥30.0 | 0.10 | ≥30.0 | ≥30.0 | 0.07 |
| D | ≥30.0 | ≥30.0 | 0.03 | ≥30.0 | 26.1 | 0.03 |
| E | 1.3 | 18.8 | 7.6 | 0.2 | 16.3 | 6.3 |
| F | ≥30.0 | 0.14 | 0.04 | ≥30.0 | 0.09 | 0.07 |
| G | ≥30.0 | ≥30.0 | ≥30.0 | ≥30.0 | ≥30.0 | ≥30.0 |
| H | 1.0 | 7.6 | 0.12 | 1.2 | 6.2 | 0.12 |
| I | ≥30.0 | 20.7 | 7.4 | ≥30.0 | 24.8 | 6.8 |
| J | ≥30.0 | ≥30.0 | 0.08 | 14.9 | ≥30.0 | 0.09 |
| K | ≥30.0 | 15.6 | 26.6 | ≥30.0 | 17.4 | 16.3 |
| L | ≥30.0 | ≥30.0 | 1.3 | 19.9 | ≥30.0 | 1.6 |
| M | 0.09 | 3.1 | 14.7 | 0.11 | 0.8 | 9.9 |
| N | ≥30.0 | 6.8 | ≥30.0 | ≥30.0 | 3.6 | ≥30.0 |
| O | 0.05 | 0.11 | 0.03 | 0.03 | 0.06 | 0.03 |
| P | ≥30.0 | ≥30.0 | ≥30.0 | ≥30.0 | ≥30.0 | ≥30.0 |

**Figure S8:**
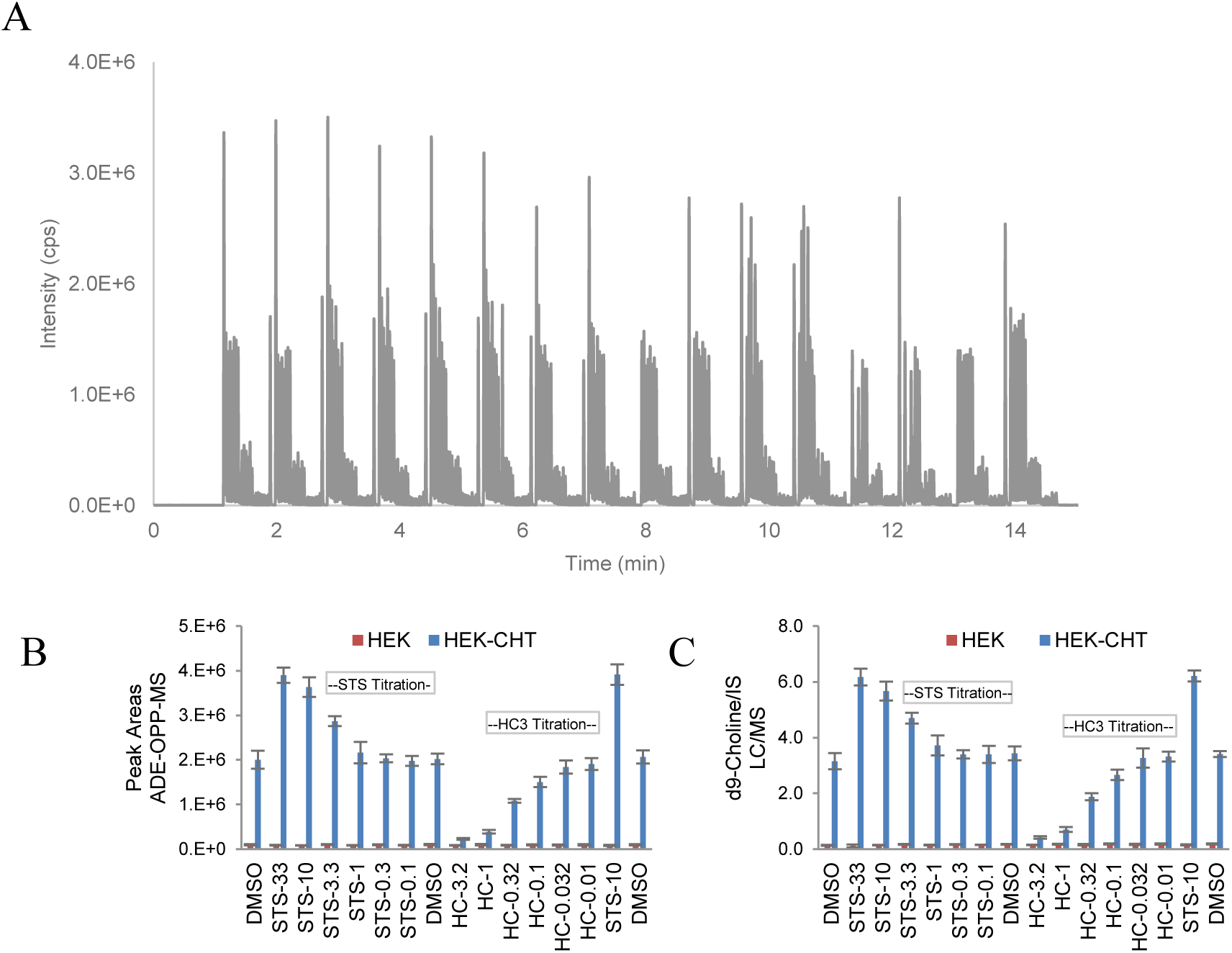
ADE-OPI-MS and LC-MS HT-Pharmacology Assay. (A) ADE-OPI-MS experimental results for a full 384-well plate for the CHT uptake assay. D9-labeled choline was monitored due to the high endogenous choline signal. All samples in the 384-wells were analyzed in ∼14 minutes. Lysed cell samples were analyzed using the ADE-OPI-MS platform (B) and compared to a conventional LC-MS method (C) (*31*). The data shown in (B) and (C) includes two modulators: hemicholinium-3 (HC), a known inhibitor and staurosporine (STS), a known activator that is dosed at a range of concentrations (*32*). On the x-axis the number after HC- or STS- represent the dose concentrations (in µM). HEK parental cell line was used as background in addition to the CHT overexpressed cell line (HEK-CHT). The ADE-OPI-MS platform increased the analysis speed 10 times and reduced sample volume 500 times while delivering equivalent data quality and concordance for both inhibition and activation profiles of the two modulators.

**Table S2:** Z’ Metrics Determined for HT-Pharmacology Assay. Shown in the table are the Z’ factors to assess the performance for 3 assays. For LC/MS, Z’ = 0.53 or 0.63 (without or with an IS). For ADE-OPI-MS, Z’ = 0.71 and there is no need for an IS. Calculation of Z’ score for inhibition CHT assay: samples with DMSO were treated as ZPEs (negative control with 0% effect), and the highest dose of HC-3 samples were used as HPEs (positive control with 100% effect).

|  | Z' | IC50 for HC3 | EC50 for STS |
| --- | --- | --- | --- |
| LC/MS without IS | 0.53 | 0.37 μM | 3.8 μM |
| LC/MS with IS | 0.63 | 0.31 μM | 3.9 μM |
| ADE-OPI-MS | 0.71 | 0.31 μM | 2.4 μM |

**Movie S1: ADE-OPI-MS system in operation.** The movie shows an ADE-OPI-MS system in operation from a 384-well microplate.

